# Energy calculations for atomic oxygen liberation from hypohalous acids biosynthesized by mammalian peroxidases

**DOI:** 10.1101/2025.03.27.645784

**Authors:** Razvan Puf, Michael L. Smith, Aatto Laaksonen

## Abstract

The ultimate process of the mammalian immunological system is destruction of invasive microbes by peroxidase (PO) activities. Three enzymes, lactoperoxidase (LPO), eosinophil peroxidase (EPO) and myeloperoxidase (MPO) require two substrates for this task. The first is H_2_O_2_ and the second either I^−^, Br^−^, Cl^−^ or SCN^−^. All POs are able to synthesize hypoiodous acid and OSCN^−^ but two can create HOBr (EPO and MPO) and only MPO can biosynthesize HOCl. Using density-functional theory (DFT) and following first principles of molecular modelling we investigated the energetics of hypohalous acid breakdowns by calculating the internal energy differences between the reactants and products. Atomic oxygen (ATOX) and simple halide acids are the energetically favoured products with halide cations and hydroxide ion being minor products. There is nothing subtle about ATOX, it is a very destructive species. ATOX production explains the documented ability of POs to indiscriminately and rapidly destroy invasive microbes and the role of MPO initiating inflammation. Previously unrecognised, the biosynthesis of ATOX also explains why mammalian POs are restricted to surfaces or special cells which are themselves destroyed after PO activation. This chemistry is the probable reason why dietary iodine, uniquely provided by the Japanese diet, helped to significantly reduce deaths from the SARS-CoV-2 viral agent during the COVID-19 pandemic in Japan.

**Highlights:**

- The energetics of hypohalous acid, HOI, HOBr and HOCl, breakdown into atomic oxygen (ATOX) or halide cation were calculated using density-functional theory
- The release of neutral ATOX from all three hypohalous acids is greatly preferred over halide cation and hydroxide production
- ATOX produced by mammalian peroxidases is incredibly powerful and largely responsible for the ultimate destruction of most invasive, especially airborne microbes

**Graphical Abstract:** 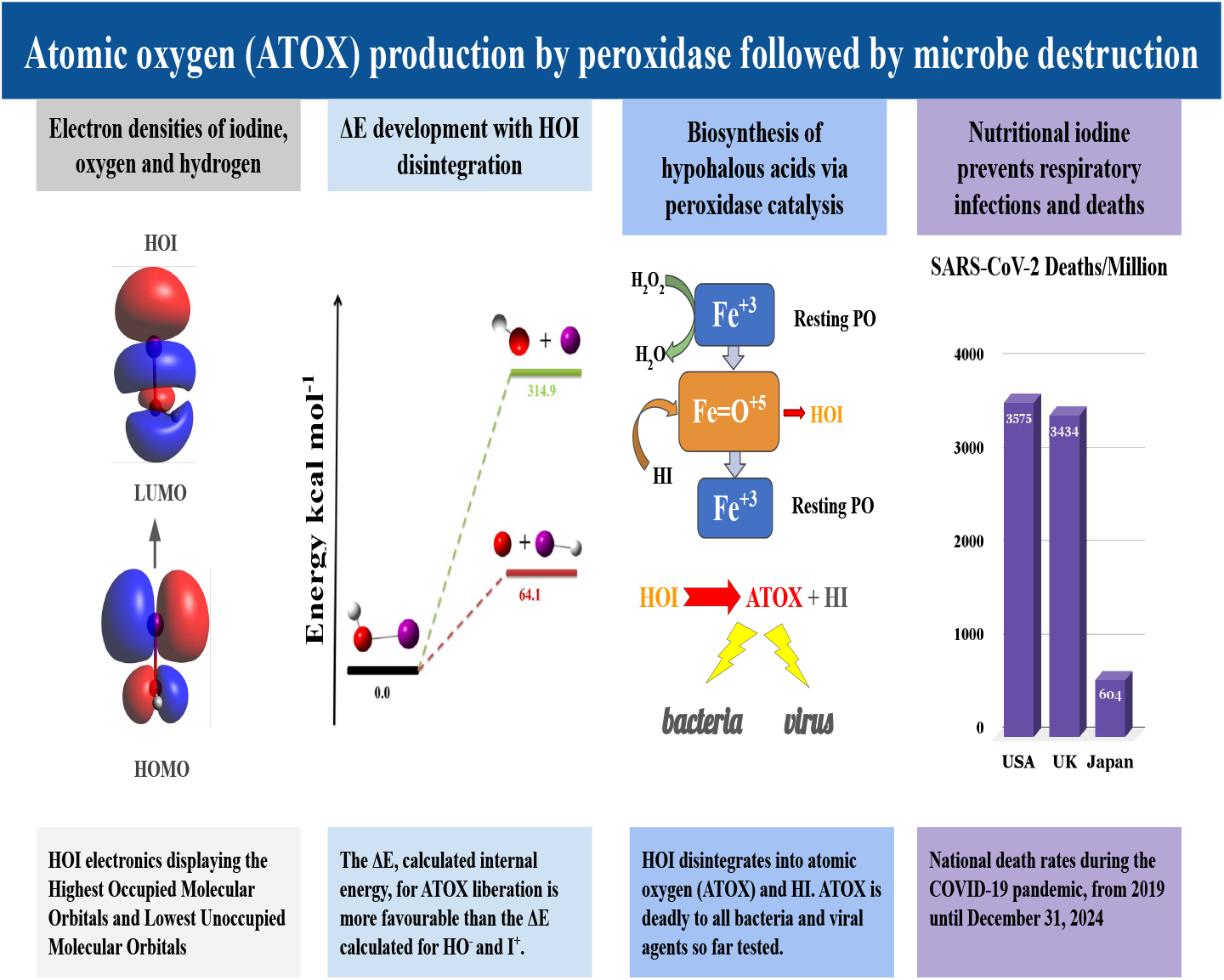

## 1. Introduction

Three mammalian peroxidases (PO, EC 1.11.1.7); lactoperoxidase (LPO) on the mucous surfaces of the nose, trachea, bronchi, eyes, mouth and in milk [14], eosinophil peroxidase (EPO) of eosinophils [31] and myeloperoxidase (MPO) abundant in neutrophils, monocytes, macrophages and natural killer (NK) cells [3]. These enzymes are responsible for the ultimate destruction of most invasive bacteria, fungi and viral agents [38]. EPO exhibits anti-parasitic and antibacterial activities and is released at sites requiring tissue remodelling [9]. MPO is a major player of innate immunity, central to the oxidative burst annihilating bacteria or viral agents engulfed by activated, protective cells while simultaneously destroying the cell itself [33]. MPO and EPO activities are also central to neutrophil extracellular trap formation leading to inflammation and morbidity [45]. The highly conserved genes for all three enzymes are located within a small region on human chromosome 17 [28] and mouse chromosome 11 [35]. The most important substrates for these POs are H_2_O_2_ and one of three halide anions; I^−^, Br^−^ and Cl^−^. The H_2_O_2_ is provided by the activity of a Duox enzyme from atmospheric dioxygen and NADPH [14].

The mechanisms of PO activities with halides are known in some detail. The reaction leading to hypohalous acids is

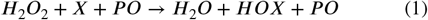

where X is I^−^ for LPO, I^−^ or Br^−^ for EPO and I^−^, Br^−^ or Cl^−^ for MPO [10]. The halide ion acts as the formal electron donor, unusual for living systems. The porphyrin iron of LPO is oxidized to the formal Fe^+5^ state during the catalytic cycle which takes just a few msec.

The pseudo-halide, SCN^−^, is also a substrate for POs, binding oxygen to become OSCN^−^ also lethal to many but not all microbes [5]. HOI has been observed at several locations around the active sight of crystalline LPO [39]. Competing reactions are the reduction of oxidised PO by a legion of biological and synthetic organic molecules oxidizing the organic and liberating water. Some of these reactions with plant peroxidases and organic dyes are commercially very important, being used for automated detection of clinical samples and many other investigative purposes.

There are three paths which hypohalous acids may follow to disintegrate in an aqueous environment. The first is the rapid, reversible proton dissociation but being low energy is not considered here. The other two are release of neutral atomic oxygen (ATOX) or production of halide cation and hydroxide. The reaction leading to ATOX production is

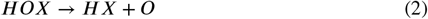

and the reaction leading to halide cation and hydroxide is

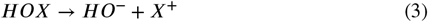

where X is either I or Br or Cl. All three hypohalous acids react with biologics producing oxidised and halogenated proteins [4]. MPO activity also modifies DNA and lipids, oxidizes and halogenates proteins [20]. A well-known, vital reaction by the related thyroid peroxidase (TPO) is the iodination of the two tyrosyl residues of the thyroid hormones, T_3_ and T_4_. Another important reaction is the crosslinking of collagen by the enzyme peroxidasin which biosynthesizes HOBr in an extracellular environment [17].

Lactoperoxidase has been shown central for maintaining health against airborne bacterial infections since resistance is impeded when LPO is removed from the airways [15]. Further studies utilizing lambs have shown that iodine offers a better defence against airborne viral agents, such as the RSV (Respiratory Syncytial Virus), than SCN^−^ [11]. It has also been demonstrated that LPO activity destroys a wide range of viral agents [26], aerobic and anaerobic bacteria [16]. The list of microorganisms; bacteria, fungi, viral agents and parasites which are liquidated by LPO has been reviewed and it is impressive [36]. The question becomes, does a microbe exist which the LPO system cannot spoil? LPO is also resistant to thermal shock over a wide pH range in response to the harsh environmental demands as LPO functions in external media [29].

Much is known about the biology of H_2_O_2_, 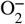, HO⋅, singlet oxygen and O_2_ which have been studied for d^2^ ecades [7, 40]. While these^2^ act to control microbes or damage living systems, none come close to the destructive powers of ATOX. It is responsible for structure decay inside nuclear reactors and low-orbit satellites [18]. The H_2_O_2_ molecule itself destroys invasive microbes but some bacteria control this oxidant by producing catalase, an enzyme which rapidly disproportionates two H_2_O_2_ into two H_2_O and O_2_ [23]. So mammals have gone a step further in self defence, developing enzymic systems producing hypohalous acids from peroxide and salts.

None of the three hypohalous acids have been isolated in pure form and may be explosive. Because of this, theoretical investigations into the energetics of hypohalous acid breakdown are warranted. Not only that but ATOX is nearly impossible to detect directly. ATOX has no magnetic moment and can only be electronically excited at very high energies or detected directly by mass spectrometry at low pressures, both conditions hostile to biology [18]. The life-time of ATOX in a biological system is probably very short. Since ATOX can only be excited by high energy ultraviolet light (UV) the signals of ATOX, as an intermediate by fast lasers, will most likely be lost in background noise. Short wavelength UV light is also likely to vaporize the biological system investigated unless experiments are performed at low temperatures. For these reasons it is necessary to investigate ATOX production using molecular modelling to estimate the initial and final state energies.

We found the three reactions liberating halide cations very energetically unfavourable but release of ATOX much less so. That all three hypohalous acids preferentially disintegrate into ATOX and simple halogen acids explains some interesting and hitherto unanswered observations. ATOX is the critical species utilised by mammals to liquidate invasive microbes which has been unrecognized in biology until now. ATOX biochemistry is probably the reason behind the recent observation that sufficient nutritional iodine intake enhances human resistance against airborne viral agents, such as the SARS-CoV-2 virus [43]. Release of ATOX is also the reason why POs are segregated to exterior surfaces and special, self-destructing cells in mammals.

## 2. Methods

We calculated the internal energies for the spontaneous release of ATOX from three hypohalous acids using first principles with density-functional theory (DFT). We also calculated the energies for release of the halide cations, I^+^, Br^+^ and Cl^+^, with simultaneous production of HO^−^. The energies, relative to the hypohalous acids, were calculated stepwise as functions of nuclear separation distances during product formation. The energy estimates were made without the presence of additional atoms, for instance, solvent.

The energies required for these bond breakages are estimated with the general relationship

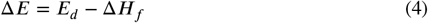

where *E*_*d*_ is the energy of dissociation and the enthalpy of formation the chemical species is Δ*H*_*f*_. Since the medium for hypohalous acid disintegration in biology is aqueous and isothermal, the pressure-volume effects are small and were neglected. Allowing this approximation we neglect the small differences expected between the internal energies and enthalpies.

For calculating the energies associated with ATOX release we use the relationships below where O* represents ATOX

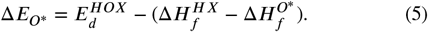

Here Δ*E*_*O*_^∗^ is the internal energy of 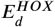 the disintegration energy of hypohalous acid, 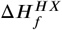 the enthalpy of halide acid formation and 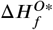 the enthalpy of formation of ATOX. For calculating the energies associated with the spontaneous halide cation release we use the relationships below where *X*^+^ represents the halide cation

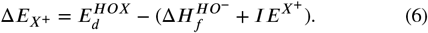

Here Δ*E*_*X*_+ is the internal energy of the halide cation, 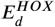 the disintegration energy of the hypohalous acid, 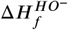 the enthalpy of formation of OH^−^ and *IE*^*X*+^ the ioniztion energy of the halide

The Density Functional Theory (DFT) method was used to determine the energies of both the reactants and products [22]. For this we relied on the B3LYP hybrid functionals with the LANL2DZ basis set [8], as well as the M06-2X functional [46] with the aug-cc-pVDZ basis set [21]. For the iodine atom, the aug-cc-pVDZ-PP basis set was employed to account for relativistic effects of this electron rich atom [27]. The halogen atom parameters used with LANL2DZ, as well as those for the aug-cc-pVDZ and aug-cc-pVDZ-PP basis sets, were extracted from the Basis Set Exchange website [32]. Calculations were performed using the Gaussian 16 software package [12]. The calculations times were quite lengthy and demanding due to inclusion of Br and especially I compounds.

## 3. Results

The calculated Hartree values for the individual species obtained after geometry optimization are listed in Table 1. The trend of increasing energy correlating with decreasing halogen atomic number is observed for all three forms; hypohaloacid, halide acid and cation halide, as expected. Also expected is the small energy difference between the hydroxide anion and the oxygen atom. These values were used to calculate the internal energies for the species listed in Table 2.

**Table 1.**
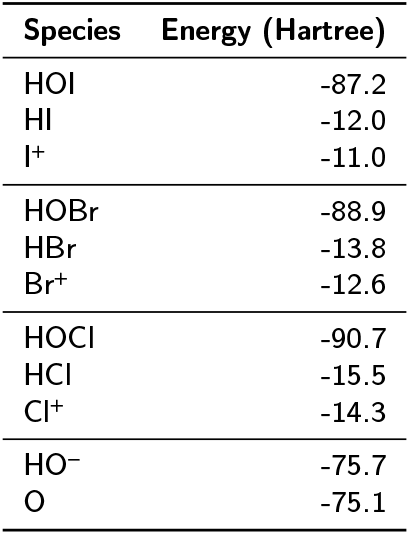
Hartree energies obtained from DFT calculations using B3LYP functionals with the LANL2DZ basis sets for reactants and products.

**Table 2.**
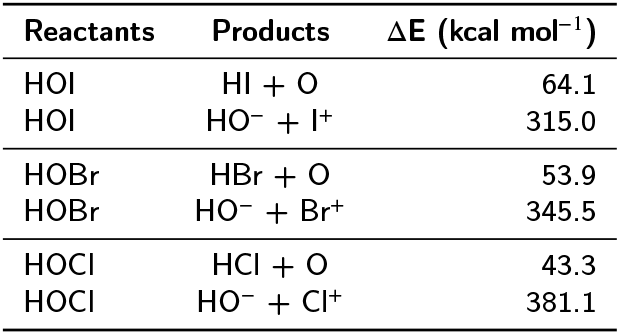
Internal energies for the disintegration of three hypohalous acids calculated from the Hartree energies of Table 1 (B3LYP functionals).

The important results are the large differences between the energies of ATOX production and those of halide cation with hydroxide productions which are presented in Table The average difference is a whopping 293 kcal mol^−1^, close to that calculated between the two paths for HOBr disintegration of 292 kcal mol^−1^. We note a correlation of decreasing energy input required to create the halide cations with decreasing halide electronegativity. That is, the creation of the ion pair requires less energy input from HOI than from HOCl, which is reassuring. Also, the release of ATOX from HOI is less favourable than release from HOBr, which is less favourable than from HOCl, again correlating with halide electronegativity. The important finding here is the great difference between disintegration energies with path, ATOX being much preferred rather than cationic halide. This is in large part due to the energy required for halide ionization along with electrical charge separation. Such large energy inputs to form the cationic halides are probably dampened, but only slightly, in the aqueous environments of biology.

**Figure 1:**
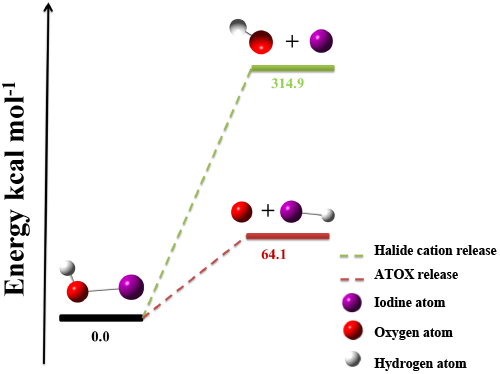
Internal energies (Δ*E*) for the disintegration of HOI, as the ground state, into ATOX + HI or HO^−^+I^+^.

## 4. Discussion

Our results demonstrate the spontaneous release of ATOX from the hypohalous acids, HOI, HOBr and HOCl is energetically favoured over halide cation and OH^−^ formations. A major reason for this large energy difference is the energies required for the electron transfer from the halide to OH along with charge separation. The liberation of ATOX is energetically favourable from all three hypohalous acids in large measure because all products are electrically neutral. While it is difficult to calculate the reaction free energies at this time we know that all three halide acids produced simultaneously with ATOX are very stable in aqueous solution and those associated free energies will be negative. Because of these reasons we think the reaction free energies in water will follow our calculated trends. We hope to improve these preliminary calculations with more detailed investigations and perhaps present the reaction free energies in the future.

There are several experimental observations consistent with our results. A straightforward experiment was performed which modelled the activity of MPO. After treatment with HOCl a target protein myoglobin (Mb) was examined by mass spectroscopy. The MW increased by steps of 16 a.m.u. but very little by 35 and 37 a.m.u. (Cl), consistent with addition of ATOX [1]. In another experiment there were indications of both ATOX and I^+^ activities when bovine and mouse serum albumins were targeted in a system utilizing MPO with I^−^ as the substrate [20]. In yet another study, the metalloproteinase, human matrilysin (MMP-7), was exposed to the action of MPO and the MMP-7 lost 4 a.m.u. This is most simply explained as two ATOXs extracting 4 H⋅ becoming two waters [13]. Our results are also consistent with the power of the LPO system with I to liquidate all microbes tested *in vitro* in several laboratories [16, 26, 36]. These experiments demonstrate that the reactions of ATOX with biologics, bacteria and viral agents are irreversible oxidation and rapid destruction.

The much smaller energy input required to create ATOX rather than an ion pair, means this reaction is preferred under the thermal conditions normal for mammals. This also means that ATOX production should be significantly enhanced when the temperature is raised by only a few degrees during the typical mammalian response to infection, if entropy allows.

Because the oxidation of organics by ATOX cannot be controlled the activities of mammalian POs are always segregated from healthy tissues. Lactoperoxidase never comes in contact with host cells, EPO and MPO after careful activation only to eliminate dangerous microbes. The results of MPO activities initiated by neutrophils, monocytes and NK cells are simultaneous microbe destruction with liquidation of these cells and nearby host cells are often damaged. Two other POs likely generating ATOX, thyroid peroxidase, requiring I, and peroxidasin, utilizing Br to cross-link extra cellular collagen, are both partitioned from most living tissues [37]. Separating POs from tissues, other than the thyroid, is obviously necessary to avoid uncontrolled tissue damage and inflammation.

The Br nutritional intake requirement is currently unknown and the human biochemistry is poorly understood. The chief source of bromine is likely table salt. It is obvious that bromine is required by eosinophils but Br may also be a mutagen if bromouracil is biosynthesized. This situation is complicated, Br required for defence and collagen biosynthesis but also creating DNA mutations, so the biochemistry and cell biology of bromine needs much more investigation. The rapid biosynthesis of hypohalous acids is often substrate limited. This is not a problem for the synthesis of HOCl by MPO where Cl^−^ is plentiful in mammals but the concentrations of Br^−^ and I^−^ can be limiting. It is estimated that over 2 billion people are iodine deficient [30]. Maintaining proper I concentration is paramount for good health but often difficult because nutritional iodine sources are rare. Indeed goitre, visible enlargement of the thyroid gland, is the classic symptom of nutritional iodine deficiency. Iodine is sparse in plant foods and highly variable in animal sources, even fish. By far the best sources of iodine are seaweed and kelp which are only commonly eaten in Japanese and some Korean cultures.

The results of our calculations for ATOX production have practical implications. The Japanese death rate during the COVID-19 pandemic was a puny fraction of that suffered by other developed societies [41]. The likely reason for this pleasant outcome is the Japanese diet which is iodine rich, encouraging liberal HOI production, therefore ATOX, by LPO in human airways [42]. We previously pointed out that the COVID-19 pandemic might be considered the largest open label study ever undertaken with iodine being the independent variable [42]. The correlation between nutritional iodine intake enhancing resistance to the SARS-CoV-2 virus, *Betacoronavirus pandemicum*, still holds today for over a half billion unknowing subjects. Another probable benefit of this iodine rich diet, from 1 to 3 mg day^−1^ adult^−1^, is the Japanese lifespan being the longest of all cultures. The Japanese people suffered the least during the COVID-19 pandemic while being the oldest of any culture. This is inversely correlated to the fact that the SARS-CoV-2 viral agent kills the oldest. This reinforces our contention that high nutritional iodine is protective.

The utilisation of iodine by all three POs explains why povidone-iodine (PVD-I) is the most widely used, effective antiseptic with very few side effects. This blend provides local POs, close to the wound, with iodine which is rapidly oxidised into HOI which becomes biocidal ATOX. A recent clinical study reported that PVD-I applied to the nasal region quickly eradicates the SARS-CoV-2 virus in human airways [6] and PVD-I application reduces SARS-CoV-2 infections [2]. This wonderful ability of PVD-I to destroy airborne viral agents has been known for many decades [24].

On the experimental side, the fundamental physics and short lifetime of ATOX makes this species very difficult to detect in the laboratory. With only electron pairs tightly held by 8 protons and 8 neutrons, ^16^O has no magnetic moment and suffers no electronic transition at energies less than high frequency ultraviolet radiation. This means that ATOX itself might only be detected in a laboratory using short burst laser radiation, in frozen samples of a PO system. Another spectral region which may be informative is detection of the rotational transitions by a microwave technique. For this it will be necessary to work with frozen samples from a PO system to dampen background radiation.

Since ATOX is so very reactive, circumstantial experimental evidence for PO generated ATOX should be rather easily uncovered. The excited triplet state of ATOX, O(3P), has been prepared in model systems and it preferentially oxidizes primary thiols such as cysteine and quickly oxidizes aryl and alkene hydrocarbons [19, 25]. The only oxygen species with an interesting magnetic signal is the expensive ^17^O. This could be incorporated into H_2_O_2_ by either glucose oxidase or a Duox enzyme and the biological products detected by NMR. Though costly, the magnetic properties of ^17^O will be highly dependent on neighbouring atoms and readily diagnostic. Both nearby ^13^C and ^1^H will be mightily affected by a neighbouring ^17^O.

Oxidation of the thioether sulphur of the common amino acid methionine is an obvious target. Methionine is a required nutrient of the human diet and has been long thought a fragile protein residue. The sulphur of methionine presents two electron pairs, just begging for attack by an electrophilic ATOX. Methionine sulfoxide is suspected of being a signal for a cell is distress [44]. As previously written, ATOX has the ability to pluck hydrogen atoms from organics. Frozen samples after ATOX exposure should display an interesting blend of free radicals which can be easily detected and characterized by EPR at low temperatures.

This report is the beginning of our investigation on the mammalian peroxidase systems and ATOX. We will continue to refine our calculations to present more exacting energies. One long-term goal is estimating the free energies of ATOX and cationic halide productions. This will demand a tremendous investment in programming and computer time. We will also work on the question of how the three peroxidase hemeproteins function in similar though slightly different manners with different halide substrates. This knowledge will help us understand why mammals have developed three different POs at three different localities.

## 5. Conclusions

Using density-functional theory (DFT) with modern functionals we have shown the major products from the breakdown of three hypohalous acids are atomic oxygen, ATOX, and halide acid with the minor products being the halide cation and HO^−^. The average difference in internal energies between these two reactions is 293 kcal mol^−1^, greatly favouring ATOX liberation from all three hypohalous acids. ATOX chemistry is simple, it oxidizes most everything biological. This is the reason why the LPO system, when supplied with enough iodine, destroys all bacterial and viral agents tested *in vivo* and *in vitro* so far. Atomic oxygen production is the primary reason why LPO, the recognized first defensive barrier against microbial infection, is too active to be located in tissues but only allowed on external surfaces, mucus and fluids. This is one of the reasons why EPO and MPO are not found in tissues but are only present in cells dedicated to immunity.

## Abbreviations

a.m.u.: atomic mass unit
ATOX: atomic oxygen, O or O*
DFT: density-functional theory
EPO: eosinophil peroxidase
EPR: electron paramagnetic resonance
LPO: lactoperoxidase
MPO: myeloperoxidase
Mb: myoglobin
NADPH: reduced nicotinamide adenine dinucleotide; phos-phate
NK: natural killer cells (innate immunity)
NMR: nuclear magnetic resonance
PVD-I: povidone-iodine
RSV: Respiratory Syncytial Virus
SARS-CoV-2: *Betacoronavirus pandemicum*
UV: ultraviolet.

## Acknowledgments

The computer resources and technical support provided by the PDC for High Performance Computing via the National Academic Infrastructure for Supercomputing in Sweden (NAISS) are gratefully acknowledged. ML Smith appreciates the help given by Niklas Hjelm-Smith in preparing the graphical abstract.

**A. Appendix**

We have also calculated the internal energies for the six disintegrations using the M06-2X functionals with the aug-cc-pVDZ basis sets. The energetics for ATOX release using this functionals and basis set are listed in Table 3. Note the trend following halogen electronegativity holds for these calculations, too. The differences between these results, which average about 13.9 kcal mol^−1^, give us an idea of the possible systematic errors for the three reactions liberating ATOX.

**Table 3.** Internal energies for the disintegration of three hypohalous acids calculated from the M06-2X functionals.

| Reactants | Products | $\Delta E \text{ (kcal mol}^{-1}\text{)}$ |
| --- | --- | --- |
| HOI | HI + O | 80.2 |
| HOBr | HBr + O | 65.7 |
| HOCl | HCl + O | 57.0 |

The internal bond strengths of hypobromous acid has been reported. The strength of the O-Br bond of the HOBr molecule was determined from the electronic spectrum in the UV region [34]. The value of 48.5 ±0.4 kcal mol^−1^ is in the neighbourhood of our 53.9 kcal mol^−1^ though these are nor estimates of exactly the same bond. This value is evidence, however, that our bond energy estimates and error about our values are close to reality.

